# Lambda N as a model substrate for studying the mechanism of Escherichia coli ATP-dependent protease Lon as a regulatory enzyme

**DOI:** 10.1101/2025.01.24.634763

**Authors:** Marianita Castro, Sanghyuk Lee, Irene Lee

## Abstract

As an ATP-dependent protease, the quality control functions of Lon have been extensively studied and reviewed in the literature. By contrast, very little research has been conducted to investigate Lon’s physiological functions and its mechanism as a regulatory protease. In this manuscript, we provided a survey of literature and data to convey that the lambda N (λN) protein is a suitable Escherichia coli Lon (ELon) substrate for studying the role played by Lon in regulating an RNA transcription process. For proof of principle, we demonstrated that the minimal component of the RNA transcription complex containing RNA polymerase (RNAP) and the σ factor can inhibit λN degradation by ELon through SDS-PAGE, and the carboxyl-terminal of λN is important for Lon competing with RNAP interaction. Using negative stain electron microscopy, we obtained structural evidence to show that λN lacking the carboxyl-terminal flanked by residues 99-107 interacted with ELon differently than full-length λN. Taken together, the activity and EM data provide a starting point for performing a physiological enzymology study on the contribution of ELon toward RNA transcription.

## 1. Introduction

ATP-dependent proteases are serine or threonine proteases containing a highly conserved ATPase module that couples the binding and hydrolysis of ATP to degrade abnormal or damaged proteins and short-lived regulatory proteins to maintain proper cellular function. ^1–7^ Based on the sequence and structural homology of the ATPase module, these proteases are classified as members of the AAA+ (**<u>A</u>**TPase-**<u>A</u>**ssociated with diverse cellular **<u>A</u>**ctivities) protein family.^8, 9^ Lon (named after E. coli bacteria that lack *lon* exhibits ELongated phenotype) and ClpA(X)P (**<u>C</u>**aseinoytic **<u>P</u>**roteinase complex partnered with subunit A or subunit X) bELong to this family. Based on the generally known functions of the ATPase module of these proteases and their propensity to degrade damaged cellular proteins, ATP-dependent proteases are considered quality-controlled protease machines. However, many of these protease machines also participate in the regulation of essential cellular processes such as nucleic acid metabolism by degrading certain short-lived timing proteins such as the lambda N (λN) antitermination transcription factor (by E. coli Lon) or the sigma subunit of RNA polymerase (RNAP; by E. coli ClpXP).^4, 10, 11^

Lon, also known as protease La, is crucial in degrading certain damaged proteins and short-lived regulatory proteins in the cells.^12–18^ This enzyme is a homo-hexameric ATP-dependent protease that resides in the cytosol of prokaryotes, lysosomes, and mitochondria of eukaryotes.^19–22, 1, 23, 24^ Mitochondrial Lon, in particular, degrades oxidatively damaged proteins, thereby maintaining mitochondrial DNA integrity and function. This function is of significant importance as it contributes to the overall health and functionality of the cell. ^25–28^ In *Saccharomyces cerevisiae*, Lon-deficient mutants suffer from large deletions in their mitochondrial genome and fail to process mitochondrial RNA transcripts.^23, 29, 30^ In some bacteria, Lon regulates the methylation of chromosomal DNA to enable proper cell differentiation^31^ or transcription of bacteriophage RNA during host infection.^32^ Understanding the fundamental mechanism of the ATP-dependent proteolytic mechanism of Lon is an essential step toward understanding the functional relationship between cellular ATP concentrations and the regulation of protein turnover events that dictate cellular functions.

## 2. Structure and function of the homo-oligomeric ATP-dependent protease Lon

ELon is a homohexamer with a molecular weight of 534 kDa ^21, 33, 34^. Oligomerization requires Mg^2+^ but not ATP^35^. Each subunit contains three functional domains: the N-terminal domain (N-domain), the ATPase domain, and the protease domain (P-domain) at the carboxyl-terminal. The N-domain is proposed to mediate substrate recognition—the Walker Box A and B motifs within the ATPase domain that bind and catalyze ATP hydrolysis. The P-domain shows the highest evolutionary conservation surrounding the proteolytic active site serine residue ^36^. Analytical ultracentrifugation and electron microscopy studies reveal that bacterial Lon proteases are hexameric ring-shaped structures containing a central cavity where the proteolytic sites reside ^34, 35^. Limited proteolytic footprinting studies of ELon reveal that adenine nucleotide protects the enzyme from nonspecific proteolysis, indicating at least one conformational change generated by binding to MgATP ^37, 38^. Insights into the structure and function of Lon are obtained from crystal structures of various forms of bacterial and human Lon ^33, 39–44^. A Ser–Lys dyad constituting the proteolytic site is also observed in the crystal structures ^33, 39, 40^. In full-length ELon, mutation of either 679Ser (the proteolytic site) or 722Lys to Ala in the catalytic dyad abolishes proteolytic but not ATPase activity ^36, 45–47^.

## 3. Functional mechanics of substrate recognition by Lon

Comparing the degradation profiles of several endogenous or heterologous substrates of ELon led to the postulate that solvent-exposed hydrophobic peptide patches in an unfolded protein function as recognition tags for the protease and initiation of substrate translocation often occurred at these tags. ^48–51^ Deleting such tags stabilizes proteins that are otherwise degraded by Lon. ^49, 51–53^ In some cases, protein degradation is initiated by incorporating a recognition tag into a stable heterologous reporter protein.^49–51^ According to the literature, other signals or features of substrates that are not hydrophobic can also target proteins for degradation by bacterial Lon. Analysis of the peptide sequences within the transcriptional activator SoxS required for Lon-mediated proteolysis shows that hydrophobicity is not necessary for forming the substrate’s recognition sequence of the substrate^51^. Studies examining the mechanism by which ELon degrades the DNA binding protein HUβ show that the recognition and binding sites within this substrate are independent of the initial cleavage site^54^. Studies with human Lon further demonstrate that the initial cleavage sites within the physiological substrate MPPα are at hydrophobic residues within structured substrate regions^55^. Gur and Sauer proposed an internal Lon recognition site known as degron in λN. ^50^ However, kinetic experiments showed that the initial binding and translocation λN occurs at the carboxyl-terminal. ^56^ Thus, the rules governing the recognition and cleavage of physiological substrates by Lon have not been clearly defined.

## 4. A working model for ATP-dependent proteolysis

A general mechanistic scheme accounting for the role of the ATPase in ATP-dependent proteolysis is illustrated in **Fig** 1. ^57^ The ATPase domain of the protease first binds/interacts with a specific recognition peptide within a protein substrate (step 1). The presence of MgATP induces a series of conformational changes within the enzyme subunits that leads to the unfolding of substrate (step 2), followed by internalization (also known as translocation) of the protein substrate into the central protease cavity (step 3). Polypeptide unfolding and translocation are proposed to constitute the rate-limiting step of the reaction ^58^and are often initiated at the recognition tag of the substrate. Both processes are hypothesized to be carried out by repetitive cycles of ATP hydrolysis that induce conformational changes within the ATP-dependent protease structure ^21, 59^ via a threading mechanism ^57^ ^60, 61^ (i.e., the polypeptide substrate is transported through the narrow central pore of the enzyme in a roughly linear conformation). In step 4, the translocated polypeptide substrates are sequestered within the central cavity of the protease machinery (the proteolytic chamber), where peptide bond cleavage occurs. ^57, 62–64^ For certain substrates of ATP-dependent proteases, proteolytic digestion generates peptides ranging from ∼ 5 to 20 amino acids but not partially digested polypeptide intermediates, concluding that degradation proceeds processively^5, 55, 57, 65^. The translocation and peptidase events can be isolated from substrate unfolding and selectively examined using unstructured substrates. Therefore, unstructured proteins such as λN, which is processively degraded by ELon in the presence of MgATP, are ideal substrates for investigating Lon’s processive protein degradation mechanism.^5, 56^ By incorporating fluorophores and corresponding fluorophore quenchers at specific sites along λN, the kinetic coordination of substrate translocation and different scissile site cleavage were determined by transient kinetic techniques to show that ELon initiates interaction with the carboxyl-terminal of λN. In the presence of MgATP, protein degradation occurs after the complete translocation of λN into the proteolytic chamber.

**Fig 1.**
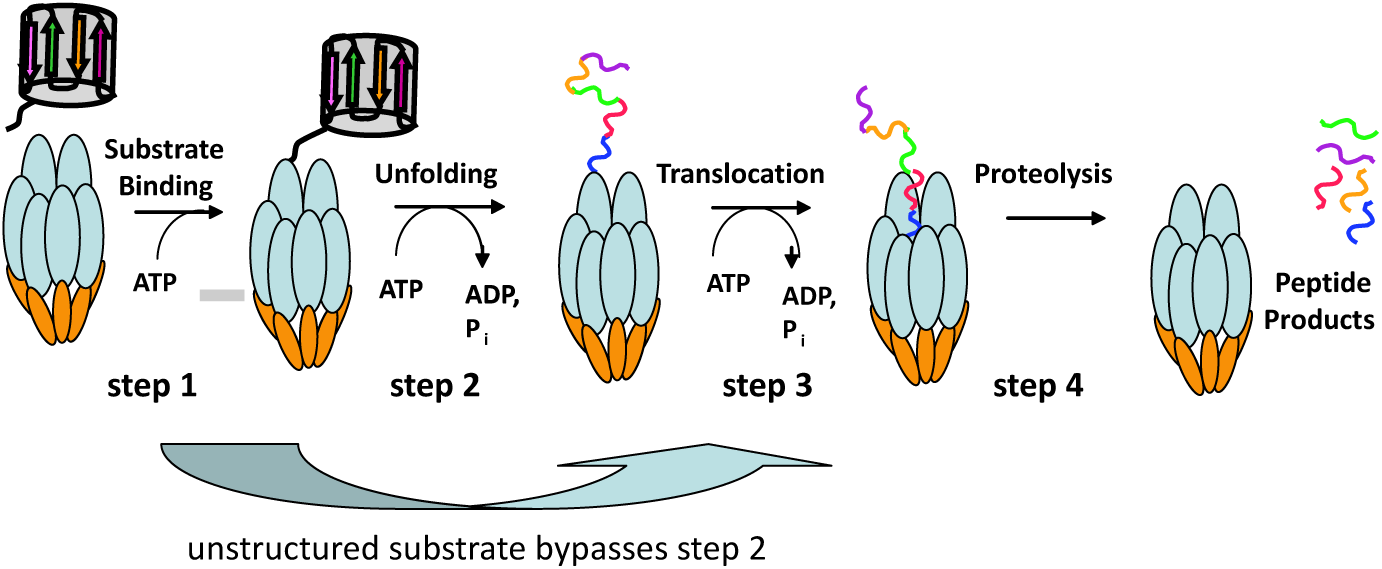
General Mechanism for ATP-Dependent Proteolysis

## 5. Lambda N as a model substrate

An intriguing aspect of the regulatory function of ATP-dependent proteases is the “timing” of substrate degradation; it is unclear how an ATP-dependent protease knows when to and not to degrade the substrate. To initiate protein degradation, an ATP-dependent protease needs to interact with at least one specific recognition element in a substrate, which may or may not be at the terminal of the substrate. Compared to the understanding of Lon functioning as a quality control protease, little is known about Lon functioning as a regulatory protease. To investigate the regulatory function of Lon, a protein substrate that functions as a timing protein must be used. To this end, we propose that the antitermination transcription protein λN is an ideal candidate.

The lambda N protein (λN) is an endogenous substrate of ELon that regulates RNA transcription during lambda phage infection of E. coli.^66, 67^ The timing of λN expression and degradation dictates whether λ phage adopts a lytic or a lysogenic life cycle in the infected host. E. coli lacking lon expression commits to lysis much faster than the wild-type cells.^68^ *In vivo*, transcriptional antitermination is processive; different regions of λ N are bound by accessory proteins and the nut RNA to form an antitermination complex. ^66, 69–71^ In vitro, λN alone can mediate non-processive transcriptional antitermination at high molar excess over the transcription complexes devoid of accessory proteins and the nut sequence.^72^ As λN interacts with the ERNAP transcription complex to carry out transcription past the termination signal in the DNA template, the impact of ELon on RNA transcription could be assessed by monitoring the kinetics of λN degradation and run-off transcription. ELon has been reported to bind to nucleic acids. In examining the proposed ELon degradation and ERNAP transcription system, one can potentially assess the impact of the nucleic acid binding affinity and selectivity of ELon towards the degradation of λN and anti-termination RNA transcription.

Nuclear magnetic resonance spectroscopic studies show that when not bound to accessory proteins or RNA, full-length and truncated forms of λN are intrinsically disordered ^67, 73–75^. Krupp et al. utilized Ala substitution and deletion mutagenesis, along with high-resolution cryo-electron microscopy, to demonstrate that R89, R96, K98/K100/K102 of λN interact with ERNAP in the transcription complex to confer antitermination activity.^76^ Although λN also interacts with ERNAP in other regions, the interactions between its carboxyl-terminal with different parts of ERNAP and nucleic acids contribute significantly to the effectiveness of the transcriptional antitermination complex. The antitermination activity of λN could be decreased stepwise by incrementally deleting the carboxyl-terminal flanking residues 85-107 but rescued by a synthetic peptide constituting residues 88-107. Work pioneered by the von Hippel and the Greenblatt labs revealed that the carboxyl-terminal of λN is important for interacting with E. coli ERNAP in the transcriptional antitermination complex.^67, 69^ Independently, NusA and boxB RNA interact with residues 34 to 47 and 2 to 19 in λN, respectively. Rees et al. demonstrated that at high concentrations (∼30-fold molar excess over transcription complexes), λN alone (without the nut sequence to supply boxB RNA and any accessory proteins such as NusA) could induce transcriptional antitermination in vitro, with an estimated K_d_ of 5 μM for the transcription complex.^77^

*In vitro*, ELon degrades purified λN in the presence of MgATP or MgAMPPNP (a non-hydrolyzable analog of ATP).^5^ Still, the rate of λN degradation is faster in the presence of ATP, and the same hydrolyzed peptide products were generated in both reactions.^5^ The cleavage profile of λN by ELon is shown in **Fig 2**. Since the unfolding of λN is not necessary, we will only need to consider the translocation and cleavage of the multiple peptide bonds within the substrate during its degradation. In vitro, full-length λN could be chemically modified or synthesized to become fluorogenic substrates for monitoring the kinetics of λN translocation and degradation at different scissile sites within the protein substrate. The K_d_ of ELon for full-length λN is 1.4 μM. ^56^ lambda N lacking the carboxyl-terminal residues 99 to 107 is less efficiently degraded by ELon.^78^ Since the K_d_ of ERNAP for λN is 5 μM whereas the K_d_ of ELon for λN is 1.4 μM, it is conceivable that Lon competes with ERNAP in binding to the carboxyl-terminal of λN. The fate of λN is dictated by the condition that favors binding to one over the other. Demonstrating that ELon can effectively compete with ERNAP in the aforementioned minimal transcription complex devoid of NUS factors and ancillary nucleic acid would lend proof of principle support for Lon’s active role in regulating RNA transcription.

**Fig 2.**
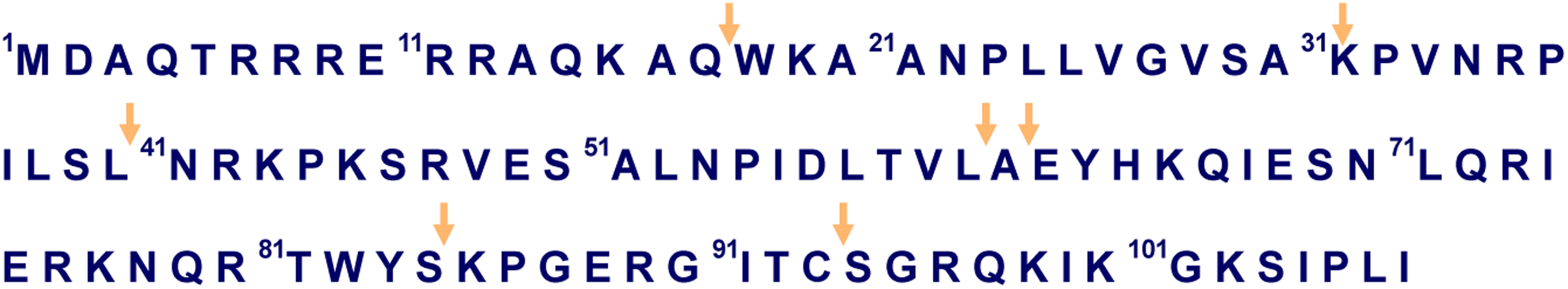
Amino Acid Sequence of lambda N. The arrows indicate the ELon cleavage sites.

## 6. A Proof of principle study

To evaluate the impact of ERNAP on ELon-mediated λN degradation, we compared the degradation profiles of full length and a truncated λN lacking the carboxyl-terminal (residues 99-107) by ELon in the presence versus the absence of the holo ERNAP (New England BioLabs), which contains the core RNAP complex and the sigma factor 70 (σ70). As noted in an earlier study, λN δ99-107 is degraded by ELon at a slower rate than full-length λN. As such, the time points for monitoring λN and λN δ99-107 are 0 to 4 min and 0 to 8 min, respectively. Each time point was quenched with SDS-loading dye and resolved in a 4-20% gradient denaturing polyacrylamide gel. Each set of assays was performed at least in triplicates. The band intensities of undigested λN proteins versus ELon were analyzed by the program Image J to produce a quantitative view of the degradation time courses. **Fig 3** shows that holo RNAP inhibits the degradation kinetics of full-length λN and λN δ99-107, but the inhibitory effect on the latter is more pronounced. This observation could be explained by ELon, but not ERNAP exhibiting weaker affinity for truncated λN and thus failing to effectively compete with ERNAP to degrade this protein.

**Fig 3.**
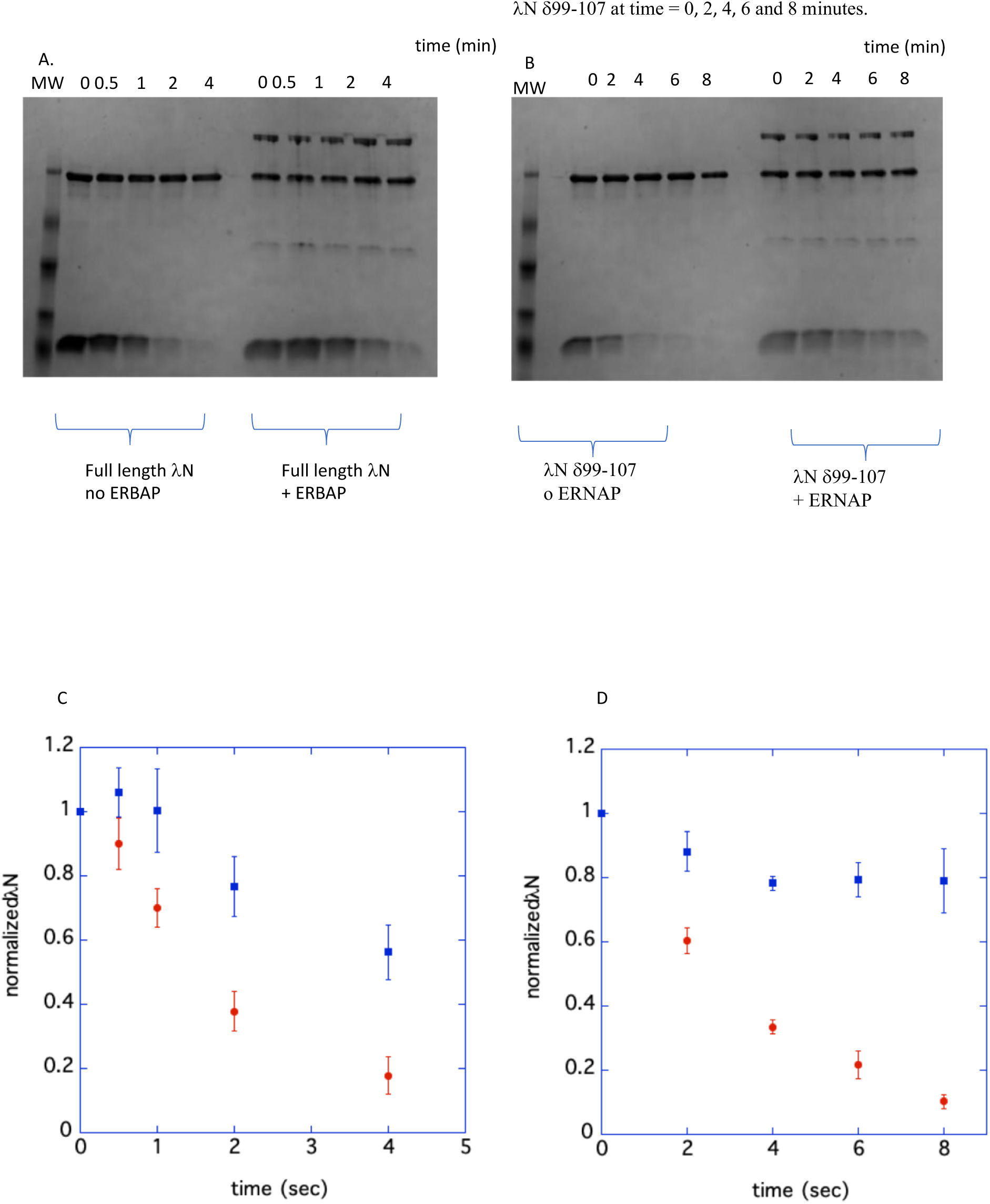
(A) A representative SDS-PAGE result of ELon degradation of full length λN at time = 0, 0.5, 1, 2 and 4 minutes. (B) A representative SDS-PAGE result of ELon degradation of λN δ99-107 at time = 0, 2, 4, 6 and 8 minutes. (C) The averaged time course of ELon degradation of full-length λN in the absence (red) versus the presence of ERNAP (blue). (D) The averaged time course of ELon degradation of λN δ99-107 in the absence (red) versus the presence of ERNAP (blue).

Negative stain electron microscopy was used to visualize the altered interactions between ELon with full length versus the carboxy-terminal truncated λN. The EM images of ELon bound to full-length or truncated λN in the presence and absence of AMP-PNP were compared. The lN proteins were not visible in the EM images, presumably due to their small size. However, structural variations were discernable in the images, indicating that λN induced structural changes in the ELon paticles. **Figs 4A to C** show the EM data on ELon by itself, whereas **Figs 4D to F** show the MgAMPPNP bound ELon. For ELon by itself, 6844 particles were selected from nine 2D class averages to produce a low-resolution 3D volume (**Figs 4B and C**). For the ELon MgAMPPNP complex, 5383 particles were selected from four 2D class averages (**Fig 4E**). The 3D volume corresponding to ELon with MgAMPPNP seems to acquire a slightly tighter conformation (**Fig 4F**). Comparing the images in Fig 4C (ELon imaged in the absence of MgAMPPNP and λN) with Fig 4F (ELon imaged in the presence of MgAMP-PNP but not λN) reveals the difference in the compactness of the particles, with the ELon: MgAMPPNP complex exhibiting a more compact structure. This finding is consistent with the limited tryptic footprinting study detecting a more compact structure in ELon when bound to MgATP or MgAMPPNP.^79^ **Figs 5A to F** show the EM results of ELon incubated with full length versus delta 99-107 λN with MgAMPPNP. To generate the 3D volumes of ELon bound to full-length and truncated λN, 7347 and 9106 particles were selected from three 2D class averages each, respectively. Comparing **Figs 4C and F** with **Figs 5C and F** reveals that ELon adopts a more compact conformation when bound to the λN proteins even without a nucleotide. A survey of the micrographs shows that the particles containing ELon bound to full-length λN in the absence of MgAMPPNP have more identifiable, uniform particles (**Fig. 5A**) than the ELon: λN δ99-107 particles (Fig 5D), presumably due to the weakened affinity of ELon for λN δ99-107 resulting in the formation of less complexes. This observation corroborates the activity findings shown in **Fig 3** that λN δ99-107 was more protected by ERNAP from ELon degradation than full-length: λN, thereby supporting the conclusion that the carboxy terminal of: λN, flanked by resides 99-107, is crucial for ELon but not so much for ERNAP interaction. **Fig 6A to C and 6D to F** show the EM data of the ELon:MgAMPPNP:: λN complexes and the ELon:MgAMPPNP: λN δ99-107 complexes, respectively. The 3D volumes of ELon: MgAMPPNP bound to full-length versus λN δ99-107 were generated from 8138 and 10020 particles selected from six and seven 2D class averages, respectively (**Figs. 6B, C, E, F**). In the presence of MgAMPPNP, the micrographs for full-length and truncated λN are similar, which may be due to more stability in the ELon: l: λN complexes gained from binding to MgAMPPNP, which promotes the translocation of the λN substrates into the proteolytic chamber (**Fig. 6C and F**). Since the full length and λN δ99-107 are degraded by ELon, albeit at a different rate (see **Fig 3**), the similarity in the EM images detected in **Fig 6** is likely attributed to the detection of ELon:λN:MgAMPPNP or ELon:λN δ99-107:MgAMPPNP translocation complex. For comparison, **Fig 7** shows the difference in the occupancy of the entrance pore of ELon bound to MgAMPPNP (Fig 7A), MgAMPPNP and λN (Fig 7B), and MgAMPPNP and λN δ99-107. The tightest hexameric conformation, which also forms propeller-like particles, is found in ELon bound to AMP-PNP and full-length lN (Fig 7B). Taken together, the EM images corroborate the proposal drawn from the λN degradation profiles summarized in Fig 3 that λN lacking the carboxyl-terminal forms a weak complex with ELon and hence is more protected by ERNAP, whose affinity for the truncated: λN remains comparable with full-length λN.

**Fig 4.**
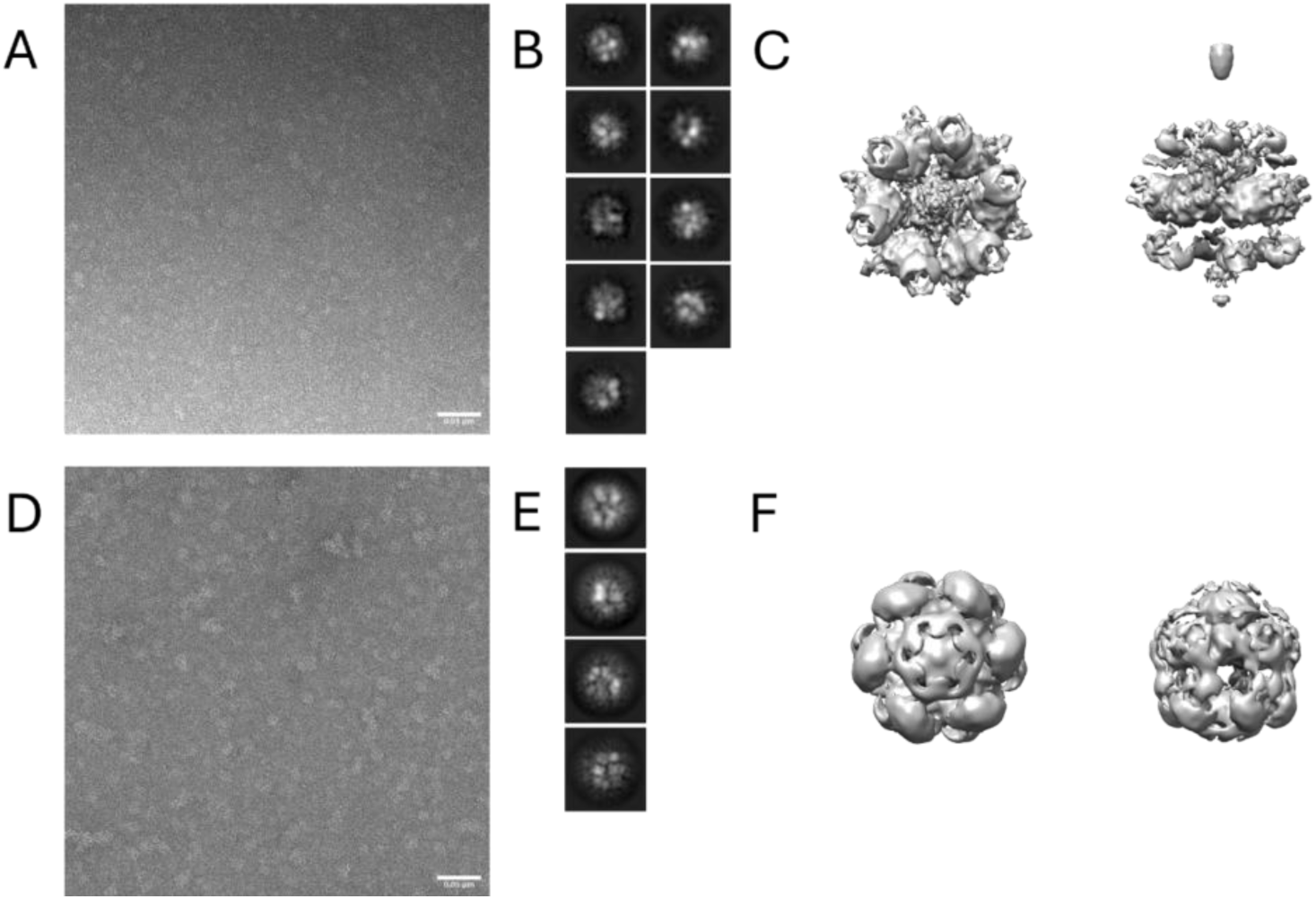
(A) Negative stain micrograph of ELon without substrate. (B) 2D class averages of ELon without substrate. (C) 3D volume of ELon without substrate with C6 symmetry. (D) Negative stain micrograph of ELon bound to AMP-PNP. (E) 2D class averages of ELon bound to AMP-PNP. (F) 3D volume of ELon bound to AMP-PNP with C6 symmetry.

**Fig 5.**
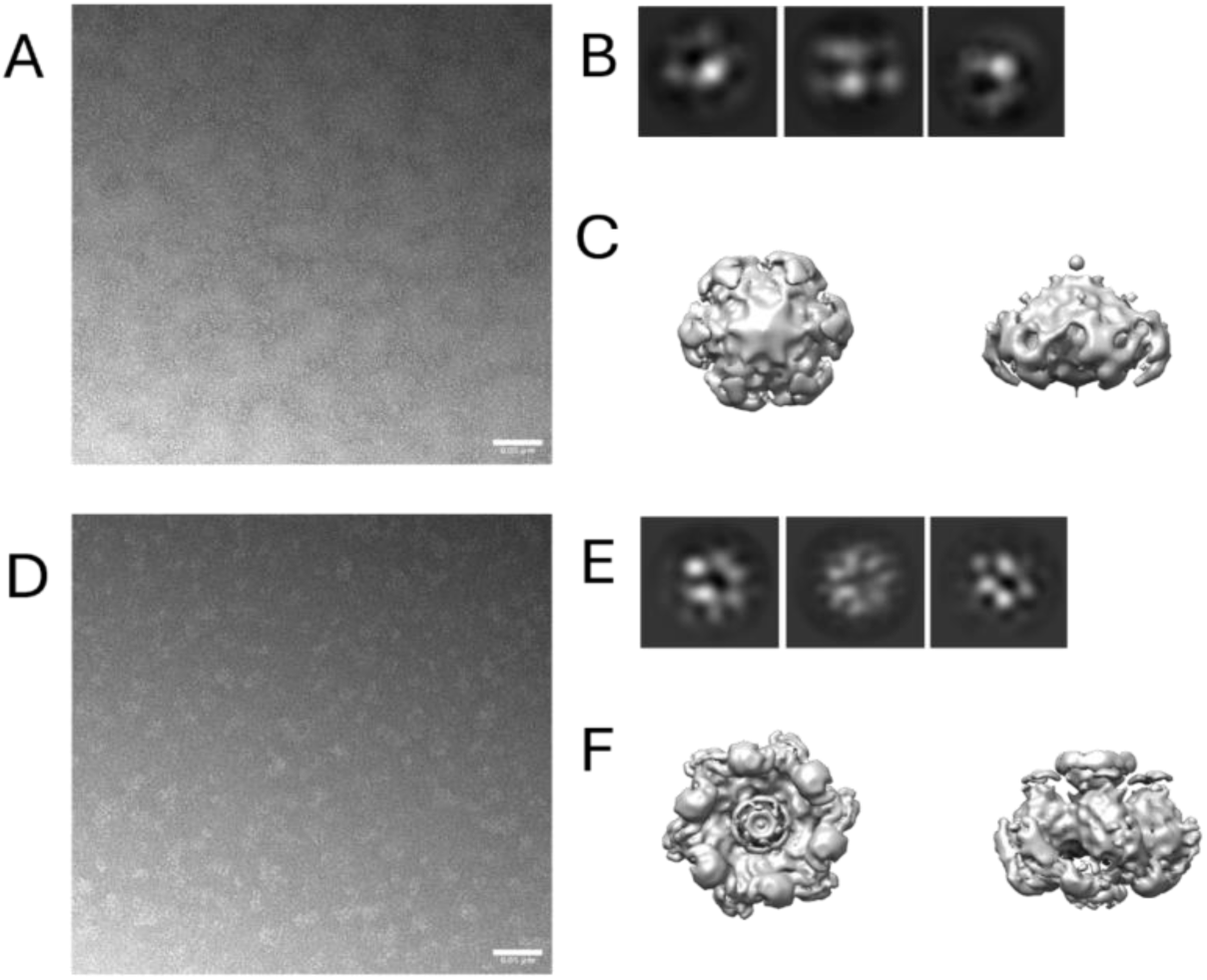
(A) Negative stain micrograph of ELon bound to full-length lambda N. (B) 2D class averages of ELon bound to full-length lambda N. (C) 3D volume of ELon bound to full-length lambda N with C6 symmetry. (D) Negative stain micrograph of ELon bound to truncated lambda N. (B) 2D class averages of ELon bound to truncated lambda N. (C) 3D volume of ELon bound to truncated lambda N with C6 symmetry.

**Fig 6.**
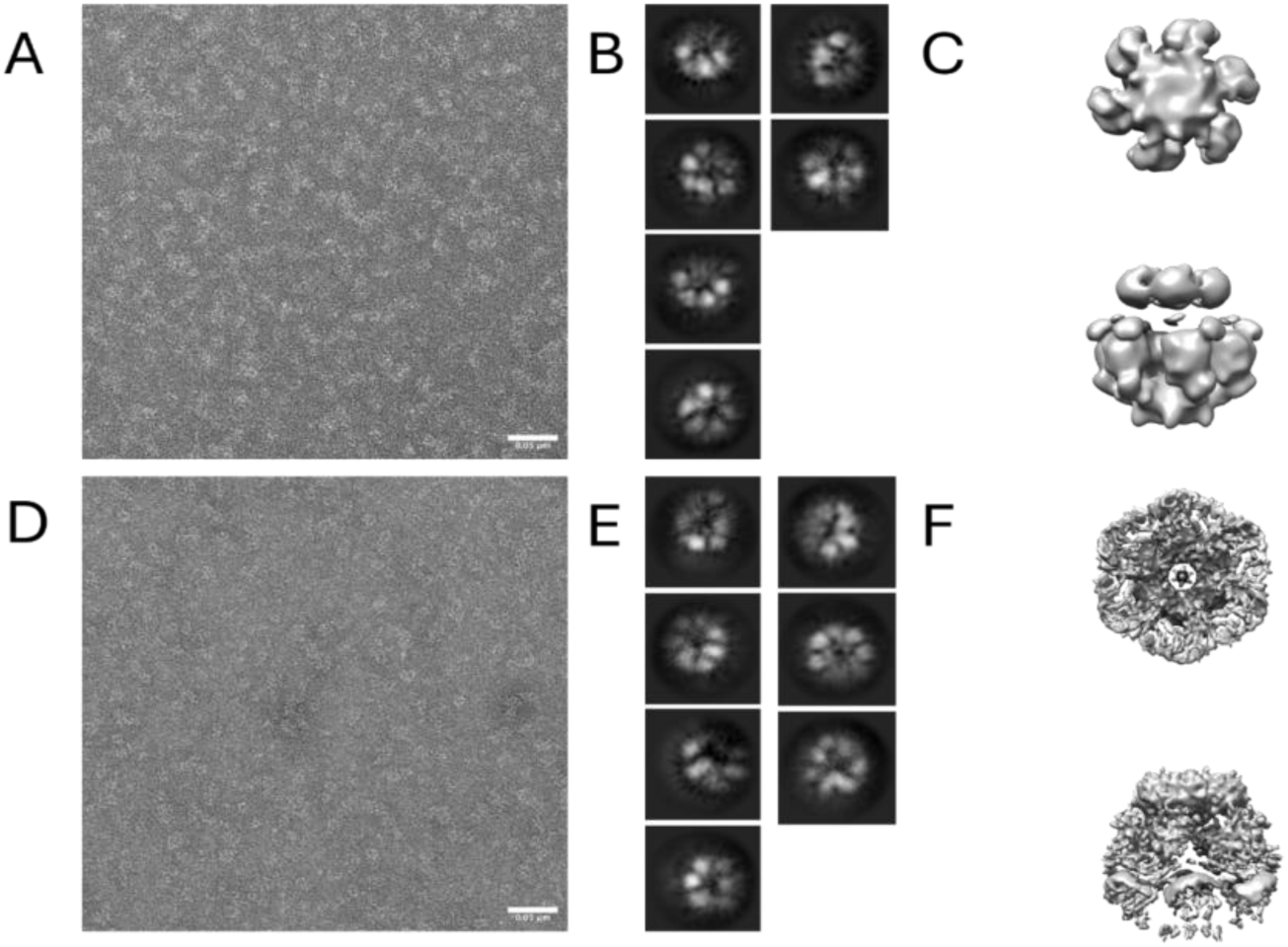
(A) Negative stain micrograph of ELon bound to full-length lambda N and AMP-PNP. (B) 2D class averages of ELon bound to full-length lambda N and AMP-PNP. (C) 3D volume of ELon bound to full-length lambda N and AMP-PNP with C6 symmetry. (D) Negative stain micrograph of ELon bound to truncated lambda N and AMP-PNP. (B) 2D class averages of ELon bound to truncated lambda N and AMP-PNP. (C) 3D volume of ELon bound to truncated lambda N and AMP-PNP with C6 symmetry.

**Fig 7.**
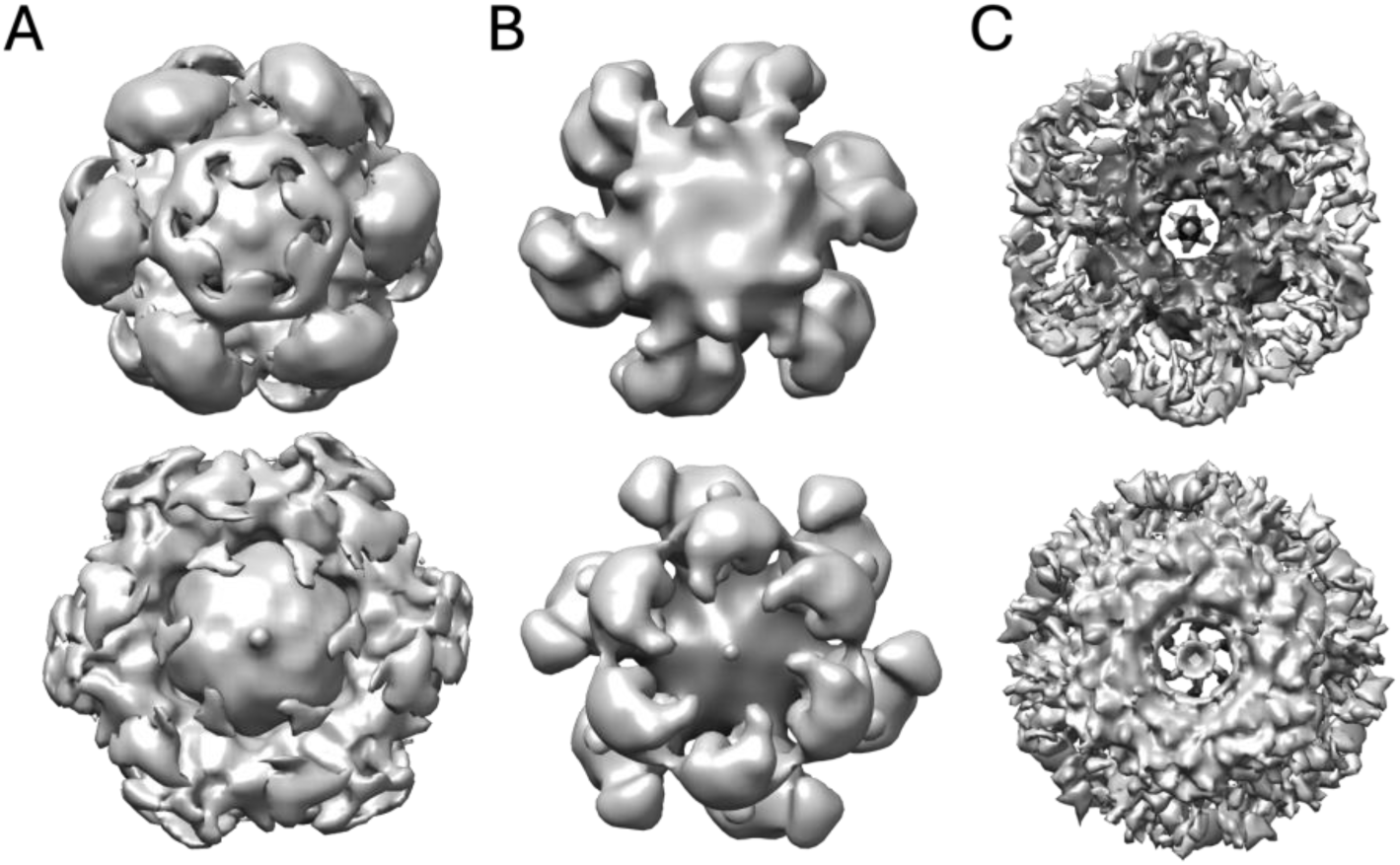
Comparison of front and back view of the entrance pore in the ELon complexes. (A) ELon bound to AMP-PNP. (B) ELon bound to AMP-PNP and full-length lambda N. (C) ELon bound to AMP-PNP and truncated lambda N.

## 7. Summary and perspective

In this manuscript, we provide literature support and primary data to support the proposal that the λN-mediated anti-termination transcription process is an excellent system for investigating the regulatory function of Lon as a protease. The contribution of λN on anti-termination RNA transcription in E. coli has been quantitatively and structurally characterized for over 30 decades; any additional structural, binding, and activity studies to evaluate the impact of ELon-mediated degradation of λN will be readily incorporated into the existing mechanism framework for RNA transcription mediated by λN. Since the E. coli RNA transcription complex constitutes the N-utilizing substance A (NusA) and the nut sequence that supplies the box B RNA in addition to the holo RNAP complex, the study of ELon mediated degradation of λN in the presence of the different RNA transcription components will provide mechanistic insights into how protein-protein interaction dictates the fate of λN, thereby affect the life cycle of lambda phage. Such knowledge will enhance our fundamental understanding of the physiological enzymology of Lon and could be applied to study the regulatory functions of other ATP-dependent proteases.

## 8. Methods and Materials

Recombinant ELon, full-length, and truncated lN proteins were prepared as described in reference 78. Chemical reagents and buffers were purchased from Sigma-Aldrich. Holo ERNAP, saturated with σ70, was purchased from New England Biolabs.

Degradation assay of full length and truncated lambda N by E. coli Lon. A reaction mixture consisting of 50 mM Tris pH 8, 5 mM Mg(OAc)_2_, 2 mM DTT, 10 μM full-length λN, 0.25 μM RNAP or RNAP dilution Buffer (NEB) was mixed with 0.5 μM ELon (monomer concentration) and 1 mM ATP. The mixture was incubated at 37°C and and time point was obtained at 0, 0.5, 1, 2, and 4 minutes by quenching an aliquot of the reaction with SDS-gel loading dye. For the assays containing λN Δ99-107, identical reagent concentrations were used, but the time points at 0, 2, 4, 6, and 8 minutes were collected. Each time point was precipitated with trichloroacetic acid and chilled at 4oC for 1 hour before centrifugation to recover the protein pellet, which was redissolved in SDS-gel loading dye loaded into a Biorad Mini-PROTEAN TGX 4-20% Precast Gel. The protein bands were detected in gel by Coomassie Blue staining and imaged using BioRad Gel Doc EZ Imager. The protein band intensities were quantified using the ImageJ software. A tif file or jpeg of the gel image was transformed into a 16-bit image type and calibrated by measuring the mean gray value of background noise under a band and then subtracting that value from the entire gel using the Math function. The lanes were selected horizontally and plotted using the Plot Lanes function to obtain intensity peaks for each band. The area under the curve was selected and measured using the Wand Tool, and the intensity ratio of lambda N to ELon was calculated using Microsoft Excel.

E. coli Lon preparation for negative stain EM and image processing. A reaction mixture of 52 mM Tris, 10 mM NaCl, 0.01 mM EDTA, 10% glycerol, 2.6 mM DTT, 1 µM ELon, 0 µM or 500 µM AMP-PNP, and 0 µM or 9.88 µM lambda N (truncated or full-length) was prepared. Samples were incubated on carbon-coated EM grids and stained with 2% uranyl acetate. Imaging was performed on a TF20 (FEI Co.) electron microscope at 200 keV. Images were captured using a Tietz 4k x 4k CCD camera at an 80,000x magnification. Particles were selected manually to generate a template picker or using a blob picker in CryoSPARC. Particles were further processed to 2D class averages. Ab-initio were generated using a C1 symmetry. Refinement was performed using a C6 symmetry since Lon typically exists as a hexamer.

## Acknowledgments

The research activities of the authors were supported by an award from the National Science Foundation MCB 2210869.

The authors wish to thank Drs. Jason A. Mears, Kristy Rochon, and Kyle Whiddon for their assistance and guidance in executing the EM imaging, data acquisition, and analyses.

## Notes

### Competing Interest Statement

The authors have declared no competing interest.

